# Differential phase partition of a PICS complex is required for piRNA processing and chromosome segregation in *C. elegans*

**DOI:** 10.1101/463919

**Authors:** Chenming Zeng, Chenchun Weng, Xiaoyang Wang, Yong-Hong Yan, Wen-Jun Li, Demin Xu, Minjie Hong, Shanhui Liao, Xuezhu Feng, Meng-Qiu Dong, Chao Xu, Shouhong Guang

## Abstract

Piwi-interacting RNAs (piRNAs) play significant roles in suppressing transposons and non-self nucleic acids, maintaining genome integrity, defending against viral infections, and are essential for fertility in a variety of organisms. In *C. elegans*, most piRNA precursors are transcribed by RNA polymerase II in the nucleus and are subjected to a number of processing and maturation steps. However, the biogenesis of piRNAs are still not fully understood. We used functional proteomics to study piRNA biogenesis in *C. elegans* and identified a PICS complex that is required for piRNA processing and chromosome segregation. The PICS complex contains two known piRNA biogenesis factors TOFU-6 and PID-1, and three new proteins PICS-1, TOST-1, and ERH-2, which exhibit dynamic localization among different subcellular compartments. In the germline of gravid animals, the PICS complex contains TOFU-6/PICS-1/ERH-2/PID-1 and largely concentrate at the perinuclear granule zone and engages in piRNA processing. During early embryogenesis, the TOFU-6/PICS-1/ERH-2/TOST-1 complex accumulates in the nucleus and play essential roles in chromosome segregation and cell division. Interestingly, the function of these factors in mediating chromosome segregation is independent of piRNA production. Therefore, we speculate that a differential phase partition of PICS factors may help cells to coordinate distinct cellular processes.

## Introduction

Piwi-interacting RNAs (piRNAs) are a class of small (21–30 bp) RNAs that associate with Piwi, a member of the highly conserved Argonaute/Piwi protein family, and play significant roles in fertility and genome stability (reviewed by ^1,2^). In *C. elegans*, piRNAs protect genome integrity in the germ line by recognizing and silencing non-self sequences such as transposons or other foreign nucleic acids, and induce chromatin modifications and neuron regeneration ^3-10^.

In *C. elegans*, piRNA precursors are transcribed from thousands of genomic loci in germline, mainly including two large genome clusters on chromosome IV ^11,12^. The <u>u</u>pstream <u>s</u>equence transcription <u>c</u>omplex (USTC), containing PRDE-1, SNPC-4, TOFU-4, and TOFU-5, has been identified to bind the Ruby motif and drive the expression of piRNA precursors ^13^. Precursors are decapped at 5’-ends to remove the first two nucleotides and trimmed at 3’-ends to remove extra-nucleotides, and generate mature piRNAs ^11, 14-16^. PARN-1, a conserved exonuclease present in germline P-granules, is required for 3’-end processing of piRNA precursors ^16^. Additionally, PID-1 and TOFU-1/2/6/7 have been found to be essential for 5’-end and 3’-end processing of piRNA precursors by forward and reverse genetic screens ^15,17^. Since most mature piRNAs are 21 nt in length and start with 5’-monophosphorylated uracil, piRNAs are also termed as 21U-RNAs in *C. elegans* ^8,11,18-20^. The term 21U-RNA is used thereafter.

21U-RNAs scan for foreign sequences, while allowing mismatched pairing with the targeted mRNAs ^7-9,21-23^. Upon targeting, the 21U-RNA/PRG-1 complex recruits RNA-dependent RNA polymerase (RdRP) to elicit the generation of secondary small interfering RNAs (siRNAs) to conduct subsequent gene silencing processes ^6-9, 21^.

However, the mechanism of how 21U-RNA precursors are processed to generate mature 21U-RNAs remains elusive. Here, using functional proteomics, we identified a <u>pi</u>RNA and <u>c</u>hromosome <u>s</u>egregation (PICS) complex that is required for piRNA processing, faithful chromosome segregation and cell division. The components of the PICS complex, TOFU-6, PICS-1, ERH-2, PID-1 and TOST-1, exhibit dynamic localization among different subcellular compartments. We propose that a differential phase partition of PICS factors may help cells to coordinate distinct cellular processes.

## Results

### TOFU-6 is required for piRNA biogenesis, chromosome segregation and cell division

A previous genome-wide RNAi screen identified seven Twenty-One-u Fouled Ups (TOFUs) that are engaged in distinct expression and processing steps of 21U-RNAs ^17^. Recently, we found that TOFU-4/5, PRDE-1 and SNPC-4 formupstream sequence transcription complex (USTC) to promote the trananscription of 21U-RNA precursors ^13^. To further understand the mechanism and roles of other TOFUs in 21U-RNA generation, we constructed GFP-3xFLAG tagged TOFU-1/2/6 transgenes (abbreviated as TOFU::GFP) using Mos1-mediated single-copy insertion (MosSCI) technology ^24^. Among them, TOFU-1/2 are expressed in the germline cytoplasm (Fig. S1A). TOFU-6::GFP is present not only in the germline cytoplasm but also at distinct perinuclear foci in the germline, which largely co-localized with the P-granule marker PGL-1 (Figs. 1A and S1B). TOFU-6 does not form granules in oocytes and embryos, which was different from PRG-1 and PGL-1 (Figs. S1B and S1C) ^19^. The TOFU-6::GFP transgene rescued the sterile phenotype of *tofu-6(ust96)*, suggesting that the transgene recapitulates the functions of the endogenous TOFU-6 protein (Fig. S1D).

**Fig. 1.**
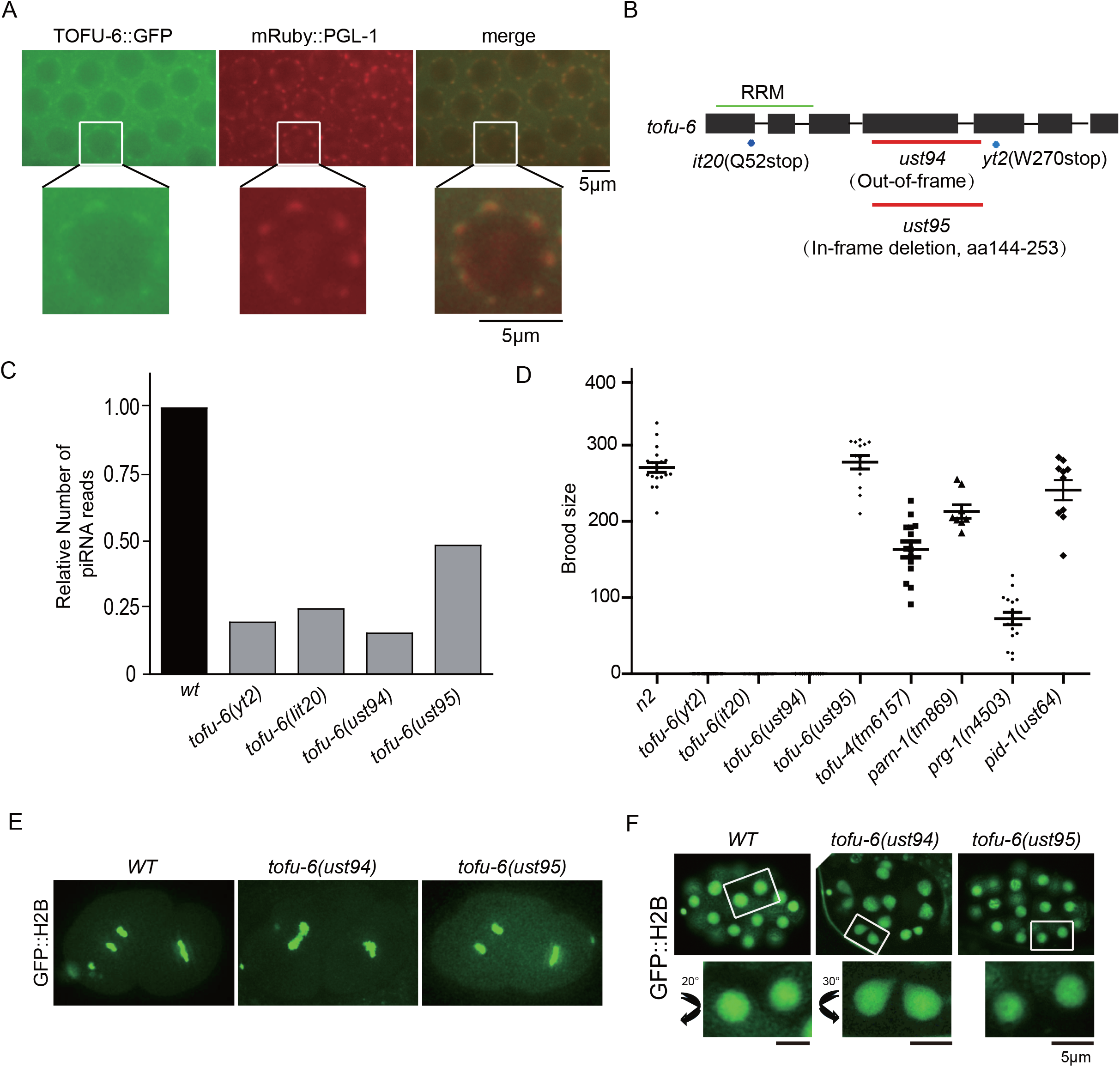
*tofu-6* is required for 21U-RNA biogenesis and chromosome segregation. (A) Images of TOFU-6::GFP and P-granule marker mRuby::PGL-1 in germ cells. (B) Schematic of four alleles of *tofu-6*. (C) Normalized 21U-RNA reads in indicated animals at late young adult stage. Indicated mutant animals were isolated from balancers. Reads were normalized to total RNA reads. (D) Bar diagram displaying brood size of indicated worms. Worms were grown at 20. (E) Images of GFP::H2B at meta-anaphase of two-cells stage embryos in indicated worms. (F) Images of GFP::H2B in indicated later stage embryos.

We focused on TOFU-6 and obtained two balanced mutants, *tofu-6(it20)* and *tofu-6(yt2)*, from Caenorhabditis Genetics Center (Fig. 1B). To facilitate genotyping and genetic manipulation of *tofu-6*, we further generated two additional deletion alleles, *ust94* and *ust95*, by dual-sgRNA-mediated CRISPR/Cas9 technology in the +/*mIn1* balancer background ^25^. The *ust94* caused a 107 amino acids deletion and frame shift, which is likely a null allele. The *ust95* caused an in-frame deletion. We collected about 1000 homozygous *tofu-6* mutant worms of each allele, and deep sequenced 18-30 nt sized small RNAs. 21U-RNAs are dramatically depleted in the *tofu-6* null mutants (Figs. 1C and S2A), which is consistent with previous results of *tofu-6* RNAi ^17^. In *tofu-6(ust95)* mutants, the amount of 21U-RNAs was reduced to roughly half of control animals. The *tofu-6(ust94)* is used as the reference allele in this work.

We then assayed the brood size of *tofu-6* mutants. *prg-1, pid-1, parn-1* and *tofu-4* mutants all have progeny although these genes are essential for 21U-RNA production (Fig. 1D) ^15-17^. Interestingly, the *it20, yt2*, and *ust94* alleles of *tofu-6* mutants are sterile while the *tofu-6(ust95)* mutant still has a brood size similar to wild type animals, suggesting that TOFU-6 has additional roles other than promoting 21U-RNA biogenesis and *tofu-6(ust95)* is defective in a part of the functions (Fig. 1D). TOFU-6 is required for the expression of PRG-1. In *tofu-6(ust94)* and *tofu-6(ust95)* mutants, the expression levels of GFP::PRG-1 were reduced (Fig S2B).

*tofu-6* mutant was previously reported with a maternal-effect lethal phenotype and was therefore named as *mel-47* ^26^. We used H2B::GFP as a chromatin marker and observed a pronounced chromosome lagging phenomenon in *tofu-6(ust94)*, but not in *tofu-6(ust95)* mutant, in early embryos (Fig. 1E). Additionally, in meta-telophase, the two daughter nuclei of *tofu-6(ust94)* were not separated completely, adopting the shape of a raindrop and with bridges between the two daughter cell nuclei (Fig. 1F).

We conclude that TOFU-6 may have dual roles in promoting piRNA biogenesis and chromosome segregation.

### Identification of TOFU-6 binding proteins by functional proteomics

We searched for proteins that interact with TOFU-6. First, we used co-immunoprecipitation followed by mass spectrometry (IP-MS) to identify candidate proteins that interact with TOFU-6 (Table S1). Next, we examined whether the identified candidates are required for the subcellular localization of TOFU-6 by feeding worms with bacteria expressing dsRNA targeting these genes. The genes that are required for the perinuclear granule localization of TOFU-6 are listed in Fig. 2A. Among them, PID-1, which functions to promote piRNA processing, was identified ^15^. In addition, we identified three new genes required for the perinuclear granule localization of TOFU-6. C35D10.13 was named as TOST-1 for twenty-<u>o</u>ne u antagoni<u>st</u>-1 (see Rodrigues and Ketting, submitted in parallel). Y23H5A.3 and F35G12.11 were named as PICS-1 (required for <u>pi</u>RNA biogenesis and <u>c</u>hromosome <u>s</u>egregation-1, see below) and ERH-2 (<u>e</u>nhancer of rudimentary <u>h</u>omolog-2), respectively.

**Fig. 2.**
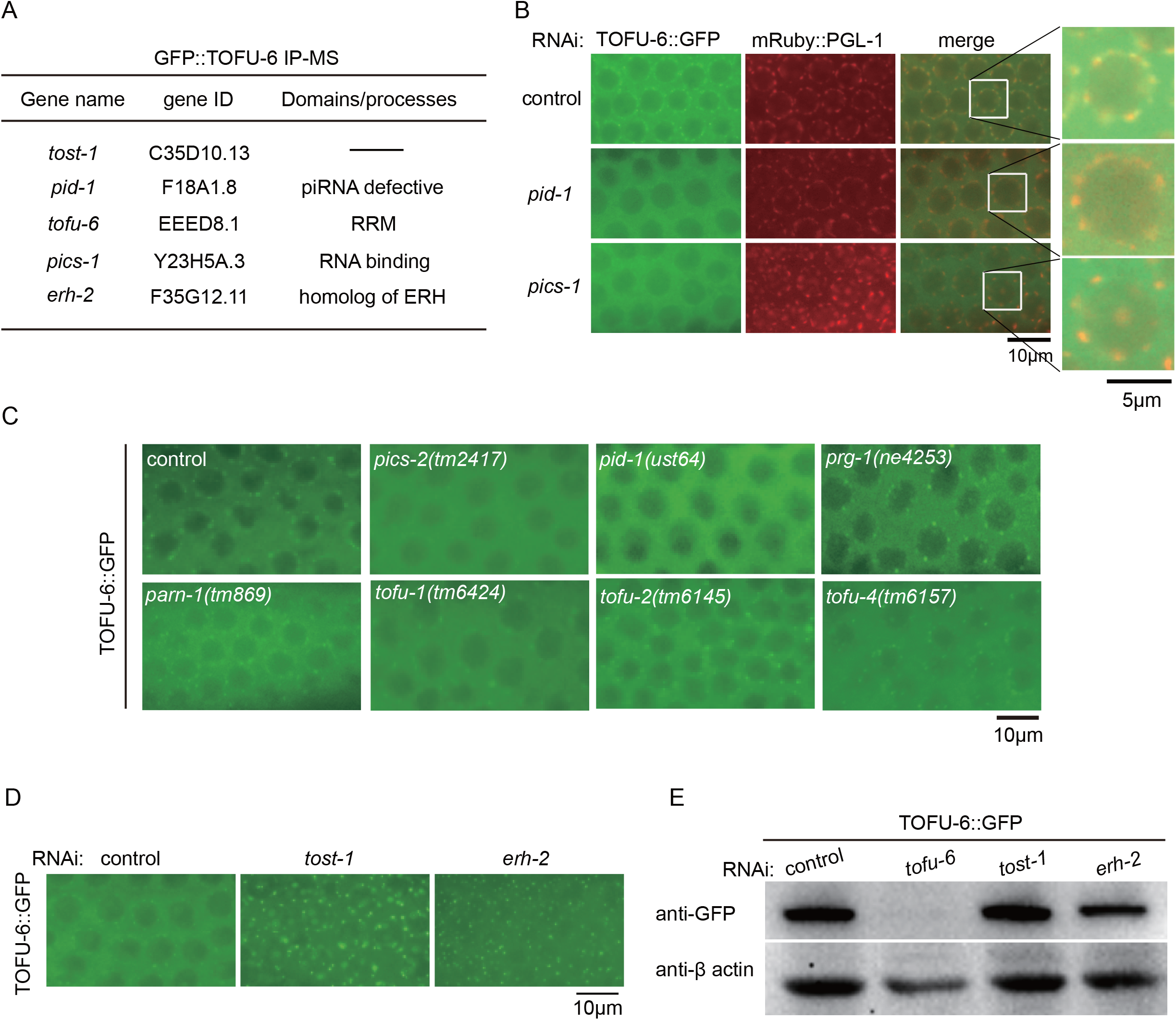
Identification of TOFU-6 binding proteins by functional proteomics. (A) Summary of IP-MS followed by feeding RNAi experiments of TOFU-6::GFP. (B) Images of TOFU-6::GFP and P-granule marker mRuby::PGL-1 after RNAi targeting *pid-1* and *pics-1*. (C) Images of TOFU-6::GFP in indicated animals. (D) Images of TOFU-6::GFP after RNAi targeting *tost-1 and erh-2*. (E) Western blotting of TOFU-6::GFP after RNAi targeting *tost-1* and *erh-2*.

When *pid-1* and *pics-1* were depleted by RNAi, TOFU-6 failed to form perinuclear granules, but the P-granule marker PGL-1 remained intact (Fig. 2B). We generated a deletion allele of *pid-1(ust64)* by dual sgRNA-mediated CRISPR/Cas9 technology and also obtained a balanced strain *pics-1(tm2417/hT2)* from National BioResource Project (NBRP) (Fig. S3A). In *pid-1(ust64)* and *pics-1(tm2417)* mutants, TOFU-6 failed to form perinuclear granules as well (Fig. 2C). The depletion of several known piRNA biogenesis factors, including t*ofu-1/2/4, prg-1* and *parn-1*, did not prevent TOFU-6 from forming perinuclear foci (Fig. 2C). However, when *tost-1* and *erh-2* were depleted by RNAi, the TOFU-6::GFP foci became bigger and brighter and exhibited mislocalization from the perinuclear granule zone (Fig. 2D). The protein levels of TOFU-6::GFP remained unchanged after RNAi *tost-1* and *erh-2* (Fig. 2E). Interestingly, in *pid-1(ust64)*, the TOFU-6 foci disassembled in meiosis region, while in *pics-1(tm2417)*, the TOFU-6 foci disassembled in both mitotic and meiosis regions (Fig. S3B).

### TOFU-6, PID-1 and PICS-1 are required for piRNA maturation

To investigate the roles of PID-1 and PICS-1 in piRNA biogenesis, we deep sequenced 18-30 nt sized small RNAs in *pid-1(ust64)* and *pics-1(tm2417)* mutants, respectively, from late-young adult worms and treated by Tabacco Decapping plus2 enzyme to remove 5′-end cap structure. 21U-RNA precursors mostly start 2 nt upstream of the 5′ end of mature 21U-RNAs and are capped with an m^7^G ^11^. Removal of 5′-end cap will facilitate the sequencing of 21U-RNA precursors. Like *tofu-6* mutants, in *pid-1* and *pics-1* mutants, the mature 21U-RNA levels were significantly reduced (Figs. 3A, S4A and S4B). In addition, in *tofu-6*, *pid-1* and *pics-1* mutants, 21U-RNA precursors were accumulated with 2 nt upstream of the 5′ end and additional nucleotides downstream of the 3′ end of mature 21U-RNAs, comparing to those of wild type animals (Fig. 3B). We conclude that TOFU-6, PID-1 and PICS-1 are involved in the processing of 21U-RNA precursors.

**Fig. 3.**
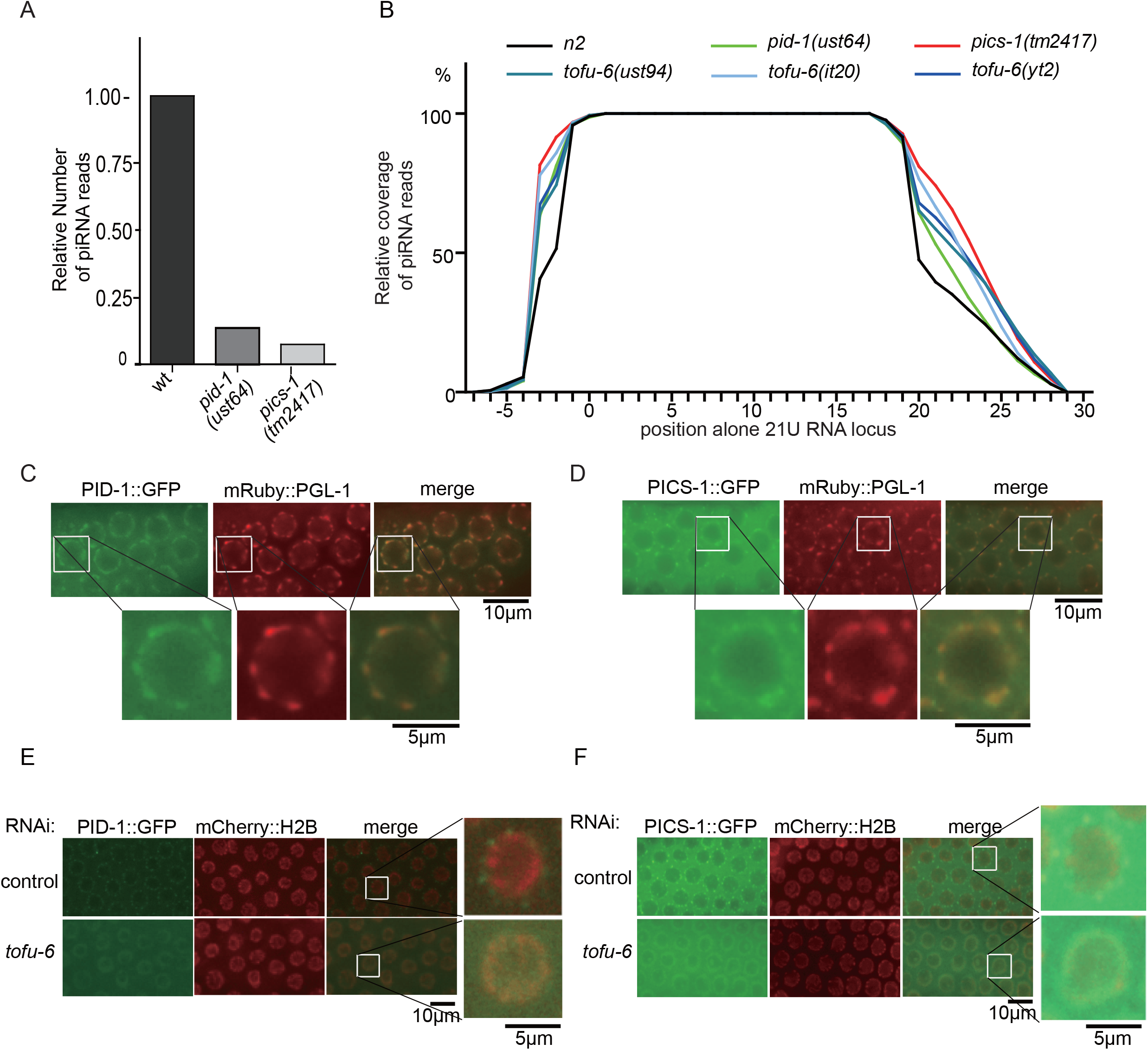
TOFU-6, PID-1 and PICS-1 are required for 21U-RNA maturation. (A) Normalized 21U-RNA counts in *N2, pid-1(ust64)* and *pics-1(tm2417)* animals at late young adult stage. *pics-1(tm2417)* worms were isolated from balancers. Reads were normalized to total RNA reads. (B) Relative coverage of individual bases of 21U-RNA loci by reads from indicated animals. The annotated 5′ ends of 21U-RNA loci are at position 0. (C, D) Images of PID-1::GFP (C) and PICS-1::GFP (D) and P-granule marker mRuby::PGL-1 in germ cells. (E, F) Images of PID-1::GFP (E) and PICS-1::GFP (F) and histone marker mCherry::H2B after RNAi targeting *tofu-6.*

We then constructed single copy GFP::3xFLAG tagged PID-1 and PICS-1 transgenes (abbreviated as PID-1::GFP and PICS-1::GFP, respectively) (Figs. S4C and S4D). The PICS-1::GFP transgene rescued the sterile phenotype of *pics-1(tm2417)* (Fig. S4E). PID-1 mainly localized in germline nuage and largely co-localized with PGL-1 as well (Fig. 3C). The expression pattern of PICS-1 was similar to that of TOFU-6, which accumulates in both germline cytoplasm and perinuclear nuage and largely overlaps with the P-granule marker PGL-1 (Fig. 3D). RNAi knocking down *tost-1* and *erh-2* induced bigger and brighter PICS-2 granules which are mislocalized from the perinuclear granule zone (Figs. S5A and S5B), similar to TOFU-6 granules after the depletion of *tost-1* and *erh-2*.

Strikingly, PICS-1::GFP and PID-1::GFP failed to form peri-nuclear granules but accumulated in the nucleus upon *tofu-6* RNAi, suggesting that PICS-1 and PID-1 may shuttle between the cytoplasm and the nucleus in the germline (Figs. 3E-F). The translocation of PICS-1 to nucleus depends on PID-1, since PICS-1::GFP failed to form perinuclear foci in meiosis region in the absence of *pid-1* (Fig. S5C) and failed to accumulate in the nucleus in *tofu-6(ust94);pid-1(RNAi)* mutants (Fig. S5D). These results are consistent with the previous report that PID-1 has both a bipartite nuclear localization signal and a nuclear export signal ^15^.

### TOFU-6, PICS-1, ERH-2, PID-1 and TOST-1 form protein complexes

To further test whether these novel factors and TOFU-6 form a protein complex, we generated single copy GFP-3xFLAG tagged TOST-1 and ERH-2 transgenes (abbreviated as TOST-1::GFP and ERH-2::GFP, respectively) (Figs. S6A and S6B). TOST-1 mainly accumulated in the cytosol of the germline syncytium, but not significantly gathered in the perinuclear granules (Fig. 4A). ERH-2 exhibits a perinuclear granule localization similar to that of TOFU-6, and largely co-localized with the PGL-1 marker as well (Fig. 4B). In the absence of *tost-1*, ERH-2 forms bigger and brighter granules yet the protein levels unaltered (Figs. S6C and S6D). Interestingly, the expression levels of TOST-1 were strongly reduced in germline and embryos but not in oocyte in the absence of *pics-1* and *erh-2* (Figs. S6E and see below Figs. 6C, S7A and S7B). Besides, ERH-2 failed to form perinuclear granules but accumulated in the nucleus when *tofu-6* and *pics-1* were depleted by RNAi (Fig. S6F).

**Fig. 4.**
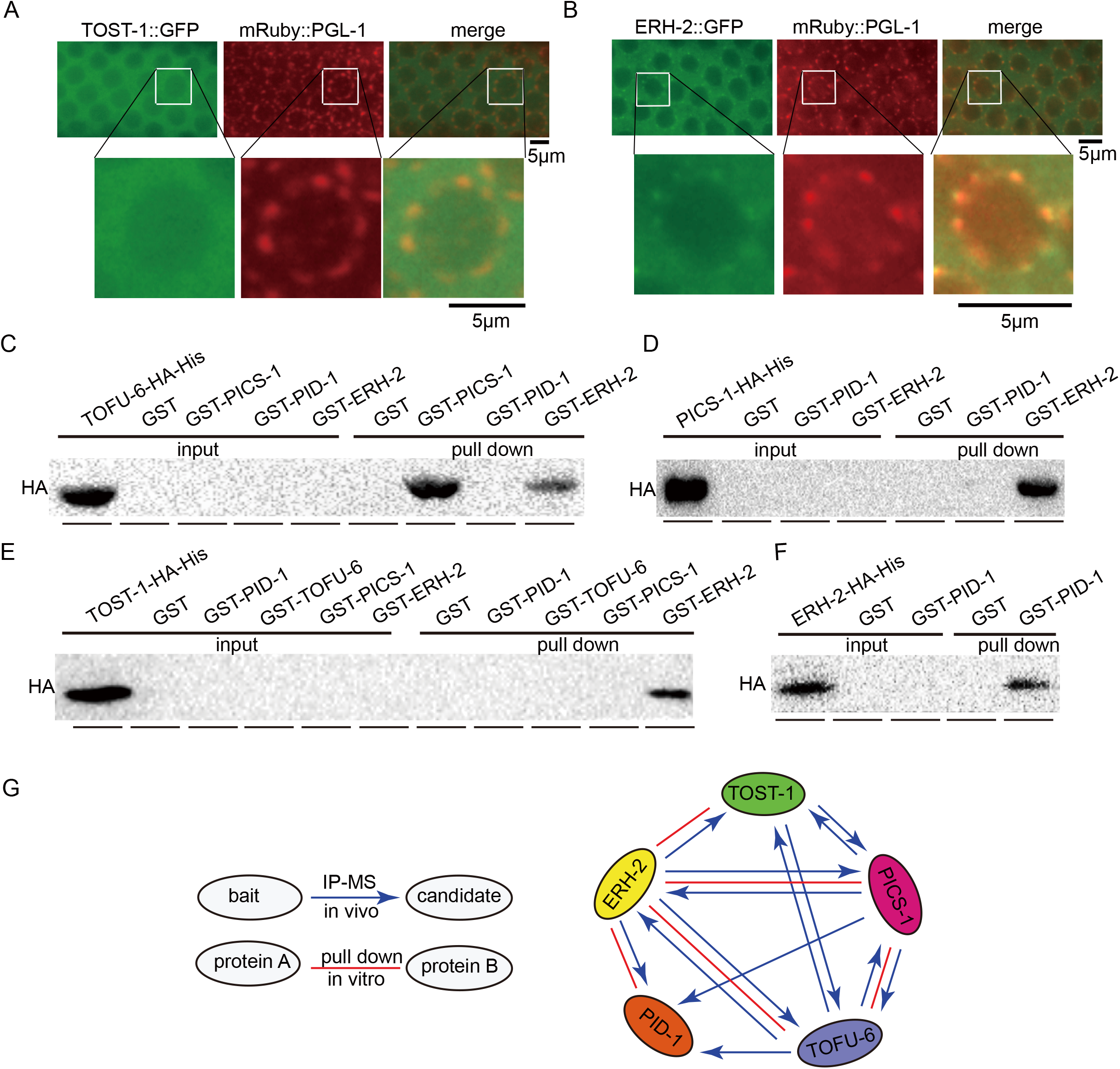
TOFU-6, PICS-1, ERH-2, PID-1 and TOST-1 interact with each other both *in vitro* and *in vivo*. (A, B) Images of TOST-1::GFP (A) and ERH-2::GFP (B) and P-granule marker mRuby::PGL-1 in germ cells. (C, D, E, F) Western blotting of pulling down samples to assay protein-protein interactions between TOFU-6, PICS-1, ERH-2, PID-1 and TOST-1 *in vitro.* (G) Summary of the protein-protein interaction of TOFU-6, PICS-1, ERH-2, PID-1 and TOST-1 both *in vitro* and *in vivo*. Red lines indicate interactions *in vitro*. Blue arrows indicate interaction *in vivo*.

We assayed the protein-protein interactions of these proteins *in vitro*. HA-His- or GST-tagged TOFU-6, PICS-1, ERH-2, PID-1 and TOST-1 proteins were expressed in *E. Coli* and purified. Then the proteins were mixed together and pulled down by GST beads. After extensive elution, the associated proteins are eluted and detected by western blot with anti-HA antibody (Figs. 4C-F). Meanwhile, we investigated the *in vivo* protein-protein interaction by immunoprecipitating 3xFLAG-GFP-tagged TOST-1, PICS-1 and ERH-2 and subjected the associated proteins by mass spectrometry (Table S1 and summarized in Figs. 4G and S7). Combining both *in vitro* and *in vivo* binding data, we concluded that TOFU-6, PICS-1, ERH-2, PID-1 and TOST-1 can form protein complexes in *C. elegans*.

Notably, although TOFU-6, PICS-1, ERH-2 and PID-1 largely co-localized with P-granules, depletion of these genes only affects the granule formation of each other (Fig. 2B), but not the formation of P-granules, as indicated by the marker PGL-1. This result suggests that the PICS factors compose distinct perinuclear granules other than P-granules per se.

### TOFU-6, PICS-1, ERH-2 and TOST-1 promote chromosome segregation and cell division

We tested whether PID-1, PICS-1, ERH-2 and TOST-1 are required for chromosome segregation and cell division, as TOFU-6 does. Using a GFP::H2B transgene, we found that TOFU-6, PICS-1, ERH-2 and TOST-1, but not PRG-1 and PID-1, are required for chromosome segregation and cell division in early embryos (Figs. 5A-C). In the absence of *tofu-6, pics-1, erh-2* and *tost-1*, but not *pid-1* and *prg-1*, we observed chromosome lagging and cell bridging phenotypes. Since PRG-1 and PID-1 are both essential for 21U-RNA production, we conclude that TOFU-6, PICS-1 and ERH-2-involved chromosome segregation and cell division processes are independent of piRNA generation. Consistently, although *pid-1* and *prg-1* mutants had viable progeny, *tofu-6* and *pics-1* were sterile (Figs. 1D and S4E). In addition, PRG-1 and PID are not expressed in oocyte and early embryos (Figs. S1B and S4D), which are different to TOFU-6 and PICS-1.

**Fig. 5.**
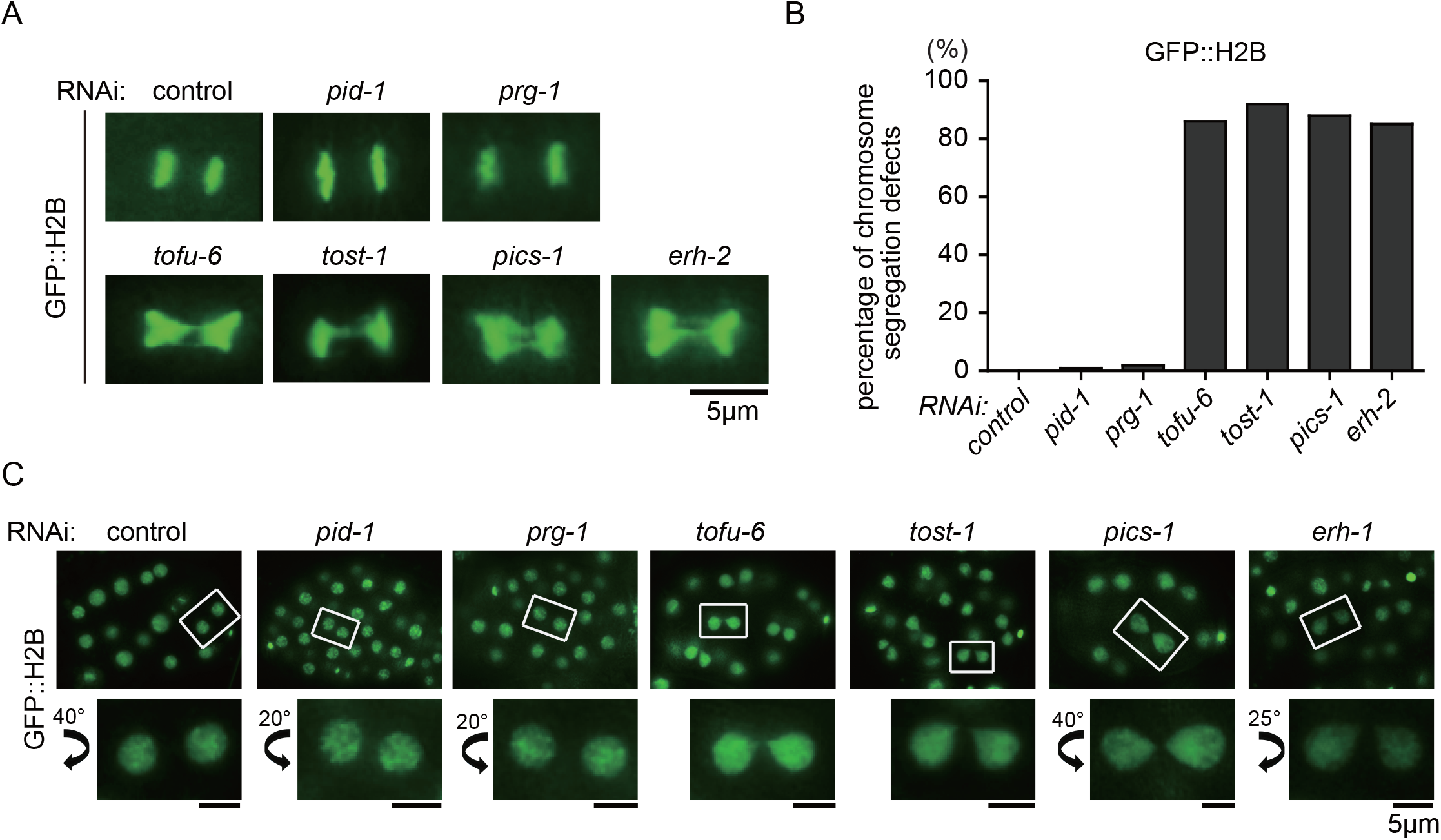
TOFU-6, PICS-1, ERH-2 and TOST-1 are required for chromosome segregation and cell division. (A) Images of GFP::H2B at meta-anaphase stage after RNAi targeting indicated genes in embryos. (B) Diagram displaying percentage of abnormal chromosome segregation. (C) Images of GFP::H2B in late embryos after RNAi targeting indicated genes.

Next, we investigated whether TOFU-6, PICS-1, ERH-2 and TOST-1 can accumulate in the nucleus. During early embryogenesis, TOFU-6, PICS-1, ERH-2 and TOST-1 enter the nucleus at the prophase of cell division, yet maintain in the cytosol at interphase without foci formation, in the 2- and 4-cell embryos (Figs. 6A and 6B). The subcellular localization of PICS factors exhibited a mutually dependent manner (Fig. 6C). For example, in all cells of early embryos, TOST-1 accumulates in the nucleus in the absence of *tofu-6*, *pics-1* and *erh-2*. ERH-2 accumulates in the nucleus in the absence of *tofu-6* and *pics-1*. PICS-1 accumulates in the nucleus in the absence of *tofu-6*. Similar nuclear accumulation of TOFU-6, PICS-1, ERH-2 and TOST-1 were observed in later embryos and oocytes (Figs. S8A and S8B).

**Fig. 6.**
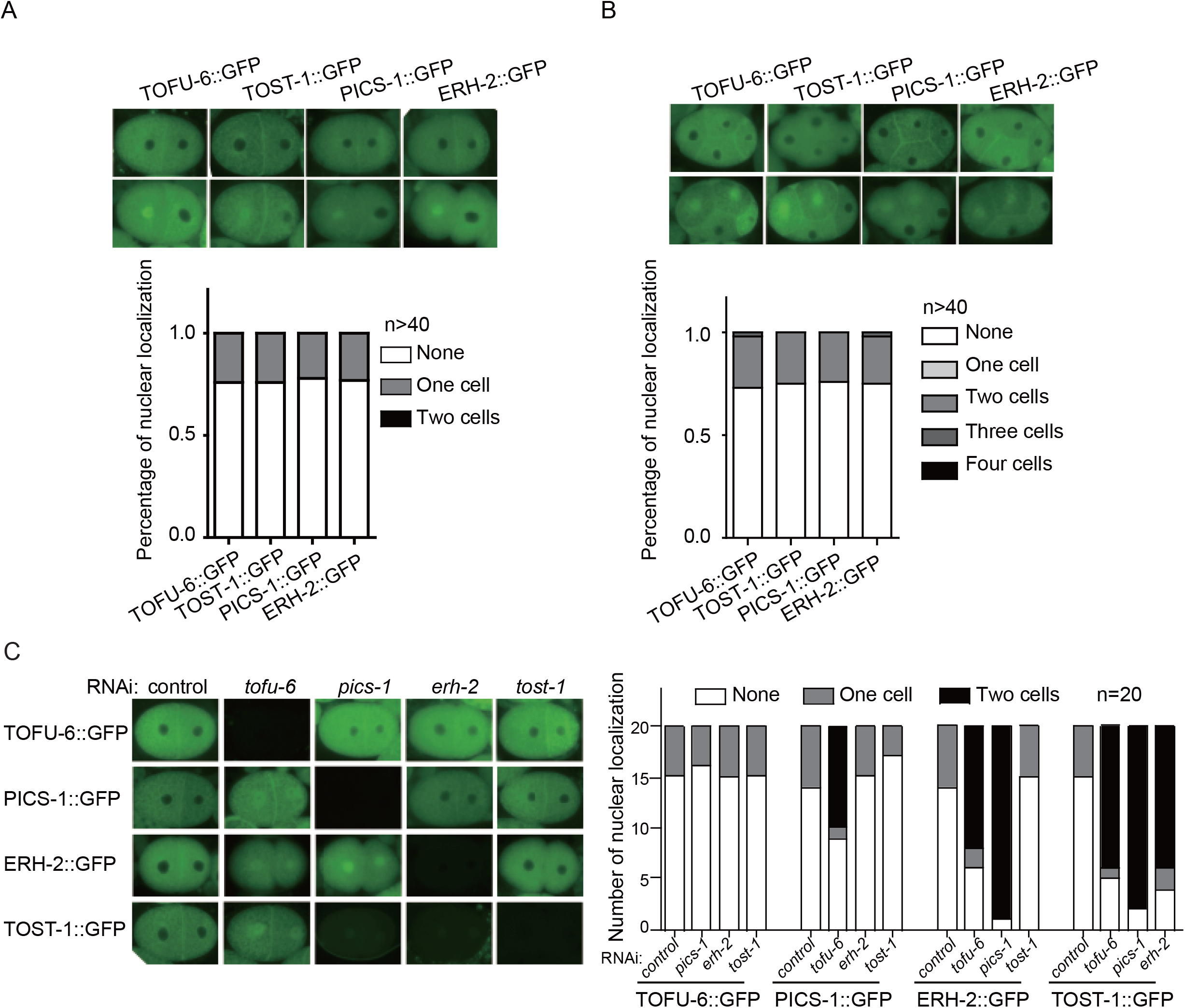
TOFU-6, PICS-1, ERH-2 and TOST-1 accumulate in nucleus during cell division. (A, B) Images of TOFU-6/PICS-1/ERH-2/TOST-1::GFP in early embryos. Diagram displaying percentage of TOFU-6/PICS-1/ERH-2/TOST-1::GFP accumulating in nucleus in two-cell-embryos (A) and four-cell-embryos (B). (C) Images of TOFU-6/PICS-1/ERH-2/TOST-1::GFP in two-cell-embryos after RNAi targeting indicated genes. Diagram displaying percentage of TOFU-6/PICS-1/ERH-2/TOST-1::GFP accumulating in nucleus after RNAi targeting indicated genes.

We conclude that TOFU-6, PICS-1, ERH-2 and TOST-1 can accumulate in the nucleus to engage in chromosome segregation and cell division processes, which is independent of piRNA biogenesis.

### Identification of IFE-3 as an additional factor required for TOFU-6 binding

To further understand the mechanism of PICS complex in mediating 21U-RNA maturation, we identified IFE-3 from the IP-MS assays of TOFU-6, PICS-1, TOST-1 and ERH-2 (Fig. S7). IFE-3 encodes one of the five *C. elegans* homologs of the mRNA cap-binding protein eIF4E, which is therefore predicted to bind capped RNAs and likely 21U-RNA precursors. *In vitro* GST pull-down assay further confirmed a direct protein-protein interaction between TOFU-6 and IFE-3 (Fig. 7A). IFE-3 largely co-localized with the P-granule marker PGL-1 in germline as well (Fig. 7B). However, the depletion of *ife-3* does not change the perinuclear localization and nuclear accumulation of TOFU-6, PICS-1, ERH-2 and TOST-1 in germlines and embryos, respectively, and vice versa (data not shown).

**Fig 7.**
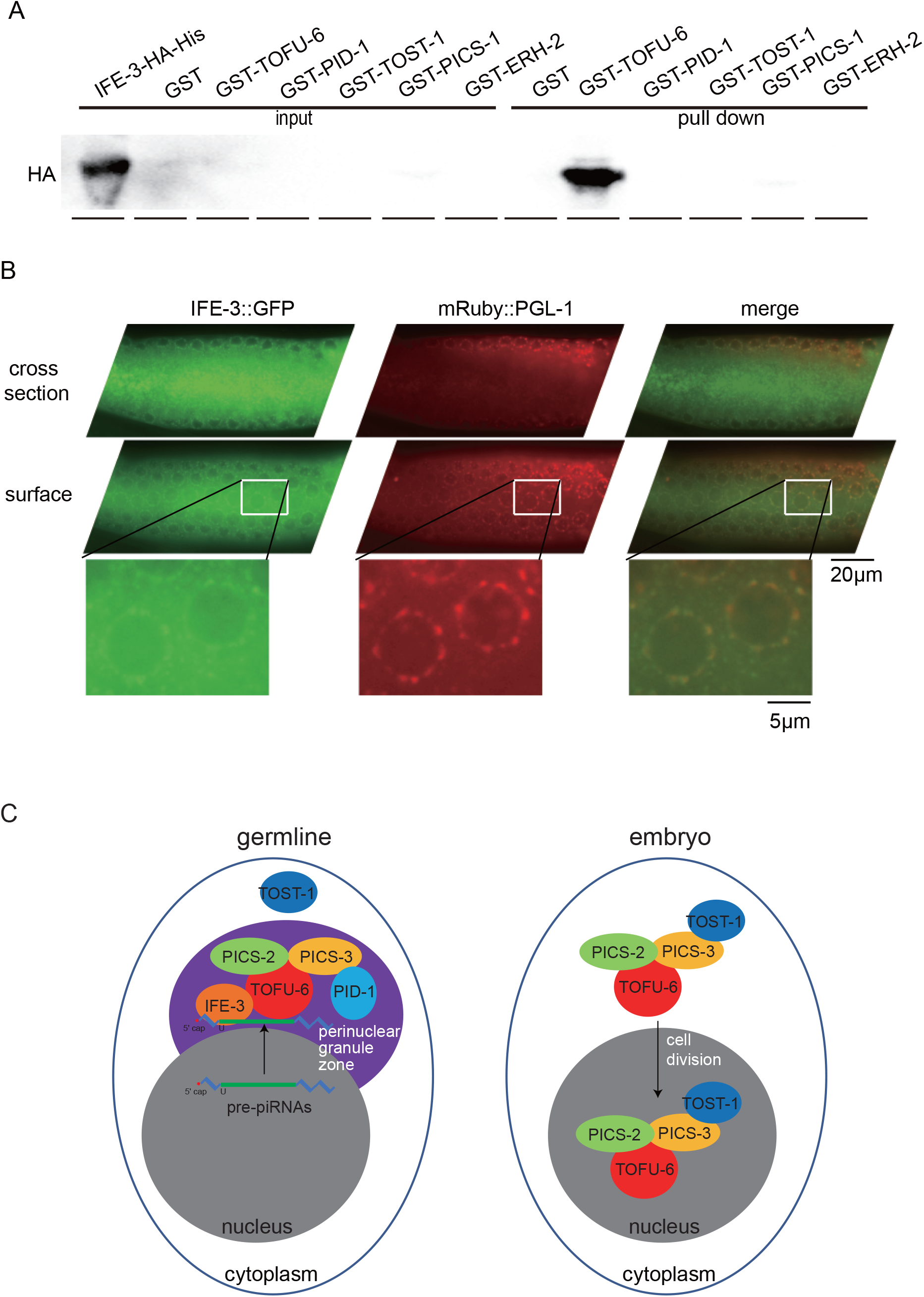
Identification of IFE-3 as an additional factor interacting with TOFU-6. (A) Western blotting of pull down samples to assay protein-protein interactions between IFE-3 and TOFU-6 *in vitro.* (B) Images of IFE-3 in germline cells. (C) A working model of the PICS complex in germline and embryos.

## Discussion

Here, by a series of proteomics and imaging experiments, we demonstrate that in the germline four proteins TOFU-6, PICS-1, ERH-2 and PID-1 might function as a complex and promote the maturation of 21U-RNAs (Fig. 7C). In embryos, TOFU-6, PICS-1, ERH-2 and TOST-1 might form another complex and accumulate in the nucleus to engage in chromosome segregation and cell division. These factors exhibit a dynamic localization among different subcellular compartments at different developmental stages. We speculate that a differential phase partition of these PICS factors may help cells to coordinate distinct cellular processes.

### PICS-1 is required for piRNA biogenesis

Previously, forward genetic screens have identified PRDE-1 and PID-1 as essential factors for piRNA biogenesis in *C. elegans* (Weick et al. 2014; de Albuquerque et al. 2014). Later, a genome-wide RNAi screening identified TOFU genes that are engaged in expression and distinct processing steps of 21U-RNAs (Goh et al. 2014). We recently found that TOFU-4/5, PRDE-1 and SNPC-4 form a USTC complex that associate with the upstream sequence element to promote the transcription of 21U-RNA precursors ^13^. Here, we combined a series of functional proteomic methods and characterized a PICS complex containing TOFU-6, PICS-1, ERH-2, and PID-1 in the germline. We further used cell biology approaches and demonstrated a mutual dependency of the components of the PICS complex to form perinuclear granules. Using deep sequencing technology, we found that 21U-RNA levels were reduced and the precursors accumulated, which suggests that the TOFU-6, PICS-1 and PID-1 proteins are engaged in the 21U-RNA maturation. However, it is unclear of the biochemical characteristics of each component of this complex. PARN-1 is a conserved exonuclease, expresses in germline P-granules, and is required for 3’-end processing of 21U-RNAs ^16^. We did not identify PARN-1 from our IP-MS experiments, suggesting that PARN-1 and PICS complex may act independent of each other to promote 21U-RNA maturation, for instance, the PICS complex may act to perform 5′ trimming of 21U-RNA precursors. Alternatively, their interaction is very transient to be captured by IP-MS experiments. Interestingly, we identified IFE-3 bound to TOFU-6, PICS-1, ERH-2 and TOST-1 from the IP-MS assay. IFE-3 binds to 5′-capped RNAs. Whether IFE-3 binds to 5′-capped 21U-RNA precursors and is involved in the maturation of 21U-RNAs requires further investigation (see Rodrigues and Ketting, submitted in parallel).

### TOFU-6, PICS-1, ERH-2 and TOST-1 engage in chromosome segregation and cell division

Strikingly, we found that *tofu-6, pics-1, erh-2* and *tost-1* mutants exhibit abnormal chromosome segregation and cell division during embryogenesis. These defects are independent of the presence of 21U-RNAs, since similar defects are not observed in *prg-1* and *pid-1* mutants. We speculate that TOFU-6, PICS-1, ERH-2 and TOST-1 may have direct roles in mediating chromosome segregation. In early embryos, TOFU-6, PICS-1, ERH-2 and TOST-1, but not PRG-1 and PID-1, could accumulate in the nucleus in mutually dependent manners.

The mechanism of how TOFU-6, PICS-1, ERH-2 and TOST-1 promote chromosome segregation is unclear. ERH (enhancer of rudimentary), the human ortholog of ERH-2, has been shown to cooperates with conserved RNA-processing factors to promote meiotic mRNA decay and facultative heterochromatin assembly, affect the replication stress response through regulation of RNA processing, and control CENP-E mRNA splicing ^27-29^. Whether TOFU-6, PICS-1 and TOST-1 act through ERH-2 to engage in chromosome segregation requires further investigation. Especially, both TOFU-6 and PICS-1 have RNA binding domains. It will be also interesting to identify the RNAs that bind to the PICS complex.

### A differential phase partition of TOFU-6, PICS-1, ERH-2, PID-1 and TOST-1 between distinct subcellular compartments

Cells organize many of their biochemical reactions in non-membrane compartments, in which proteins and nucleic acids are concentrated. Recent evidence show that many of these compartments are liquids that form by phase separation from the cytoplasm, including many PZM-granule factors in *C. elegans* ^30^. These condensates are involved in diverse processes, including RNA metabolism, ribosome biogenesis, the DNA damage response and signal transduction ^31^. We show here that certain piRNA processing factors, including TOFU-6, PICS-1 and ERH-2 localize to the perinuclear condensate which we named the PICS granules. We found that these factors were distributed in the cytoplasm, the perinuclear nuage, and the nucleus in mutually dependent manners. Moreover, the PICS granules exhibit a spatiotemporal distribution at distinct developmental stages. While they assembled as perinuclear nuage in the adult germline, they disassembled in oocytes and early embryos. We speculate that the differential phase partition of this liquid-like condensates in different subcellular compartments helps to organize and coordinate distinct cellular processes, including piRNA biogenesis, chromosome segregation and cell division. Whether other gene regulatory or biochemical pathways can be organized and coordinated in similar ways needs further investigation.

In adult germ cells, PZM granules contain ordered tri-condensate assemblages with Z granules, P granules and Mutator foci ^30^. Here, we identified PICS granules largely co-localized with the P-granule marker PGL-1. We speculate that there are a number of distinct liquid droplet organelles localized in the perinuclear granule zone (PGZ), ordered in a temporal and spatial fashion, and therefore organizing and coordinating the complex RNA processing pathways that underlie gene-regulatory systems. These liquid droplet organelles may exchange their protein and RNA components with each other and other nuclear and cytoplasmic compartments. With emerging high resolution imaging technologies, it will be great to dissect the compositions, dynamics and biological roles of this perinuclear granule zone.

## Materials and methods

### Strains

Bristol strain N2 was used as the standard wild type strain. All strains were grown at 20°C unless specified. The strains used in this study were listed in Supplementary Table S2.

### Construction of transgenic mutant strains

For TOFU-1::GFP, TOFU-2::GFP, TOFU-6::GFP, PID-1::GFP, PICS-1::GFP, TOST-1::GFP, endogenous promoter sequences, 3’ UTR, ORFs, coding sequence of gfp::3xflag and a linker sequence (GGAGGTGGAGGTGGAGCT) (inserted between ORFs and gfp::3xflag) were fused and cloned into PCFJ151 vectors using ClonExpress^®^ MultiS One Step Cloning Kit (Vazyme C113-02, Nanjing). These transgenes were integrated onto the *C. elegans’* chromosome II by MosSCI technology.

For ERH-2::GFP, IFE-3::GFP and GFP::PRG-1, coding sequence of gfp::3xflag was inserted before stop code or after initiation code using CRISPR-Cas9 system. Repair templates contained homologue arms of 1000bp to 1500bp and were cloned into vector using ClonExpress^®^ MultiS One Step Cloning Kit (Vazyme C113-02, Nanjing). Injection mix contains PDD162 (50 ng/μL), repair plasmid (50 ng/μL), pcfj90 or 90-gfp (20 ng/μL) and one or two gRNAs close to N-terminal or C-terminal of the genes (20 ng/μL). Mix was injected into young adult N2 animals. Two or three days later, F1 worms containing pcfj90 or 90-gfp marker were isolated. After growing at 20°C for another three days, animals were screening for GFP insertion by PCR.

For gene deletions, 3 or 4 sgRNAs were co-injected into N2 or +/mIn-1 animals with PDD162 (50 ng/μL), 90-gfp (20 ng/μL), 10xTE buffer, DNA ladder (500 ng/μL). Two or three days later, F1 worms expressing 90-gfp marker were isolated. After growing at 20°C for another three days, animals were screened for deletion by PCR.

Primers used for molecular cloning and dual-sgRNA-directed CRISPR/Cas9-mediated gene deletion are listed in Supplementary Table S3.

### Immunoprecipitation followed by mass spectrometry analysis

The mix-staged transgenic worms expressing TOFU-6::GFP, TOST-1::GFP, PICS-1::GFP, and ERH-2::GFP were resuspended in the same volume of 2x lysis buffer (50 mM Tris-HCl pH 8.0, 300 mM NaCl, 10% glycerol, 1% Triton-X100, Roche®cOmplete™ EDTA-free Protease Inhibitor Cocktail, 10 mM NaF, 2 mM Na_3_VO4) and lysed in a FastPrep-24™ 5G Homogenizer. The supernatant of lysate was incubated with home-made anti-GFP beads for one hour at 4°C. The beads were then washed three times with cold lysis buffer. GFP immunoprecipitates were eluted by chilled elution buffer (100 mM Glycine-HCl pH 2.5). About 1/8 of the eluates were subjected to the western blotting analysis. The rest of the eluates were precipitated with TCA or cold acetone and dissolved in 100 mM Tris, pH 8.5, 8 M urea. The proteins were reduced with TCEP, alkylated with IAA, and finally digested with trypsin at 37°C overnight. The LC-MS/MS analysis of the resulting peptides and the MS data processing approaches were conducted as previously described ^32^.

### RNAi

RNAi experiments were conducted as previously described ^33^. Images were collected using a Leica DM2500 microscope.

### RNA isolation and sequencing

Synchronized late young adult worms were sonicated in sonication buffer (20 mM Tris-HCl [PH 7.5], 200 mM NaCl, 2.5 mM MgCl_2_, and 0.5% NP40). The eluates were incubated with TRIzol reagent followed by isopropanol precipitation. The precipitated RNAs were treated with calf intestinal alkaline phosphatase (CIAP, Invitrogen), re-extracted with TRIzol, and treated with Tabacco Decapping plus 2 (Enzymax).

Small RNAs were subjected to deep sequencing using an Illumina platform (Novogene Bioinformatics Technology Co., Ltd). Briefly, small RNAs ranging from 18 to 30 nt were gel-purified and ligated to a 3’ adaptor (5’-pUCGUAUGCCGUCUUCUGCUUGidT-3’; p, phosphate; idT, inverted deoxythymidine) and a 5’ adaptor (5’-GUUCAGAGUUCUACAGUCCGACGAUC-3’). The ligation products were gel-purified, reverse transcribed, and amplified using Illumina’s sRNA primer set (5’-CAAGCAGAAGACGGCATACGA-3’; 5’-AATGATACGGCGACCACCGA-3’). The samples were then sequenced using an Illumina Hiseq platform.

### Bioinformatics

The Illumina-generated raw reads were first filtered to remove adaptors, low-quality tags and contaminants to obtain clean reads at Novogene. Clean reads ranging from 18 to 30 nt were mapped to the unmasked *C. elegans* genome and the transcriptome assembly WS243, respectively, using Bowtie2 with default parameters. The number of reads targeting each transcript was counted using custom Perl scripts.

### Brood size

Ten L3 worms were singled to fresh NGM plates and the number of progenies were scored.

### Protein expression and purification

The GST-tagged proteins were cloned into the vector PGEX-4T-1 and HA-tagged proteins were cloned into the vector PET22b. All expression plasmids were transformed into *E. coli* Rosetta(DE3) and proteins were expressed at 16 for 20 hours in the presence of 0.5 mM IPTG. The recombinant proteins, which contain a N-terminal GST-tag, were purified on a Glutathione sepharose (GE Heathcare). And the recombinant proteins, which contain a C-terminal HA-tag and 6 ×His tag, were purified on a Ni-NTA resin (GE Heathcare).

### GST pull-down

Recombinant GST-fused proteins were incubated with Glutathione sepharose (GE Healthcare) in buffer (20 mM Tris, 200 mM NaCl, PH 7.5) for 40 min at 4. Then HA-tagged protein were added and incubated 40 min at 4. The resin was then washed three times with buffer (20 mM Tris, 200 mM NaCl, PH 7.5, 2 mM DTT, 0.1% NP-40). Finally, 50 mM glutathione was added and incubated 30 min to elute the protein. All GST pull down samples were loaded on SDS-PAGE and then transfer to Hybond-ECL membrane for western blotting.

### Western Blotting

Proteins were resolved by SDS-PAGE on gradient gels (10% separation gel, 5% spacer gel) and transferred to Hybond-ECL membrane. After washing with TBST buffer (Sangon biotech, Shanghai) and blocking with 5% milk-TBST. The membrane was incubated at room temperature for two hours with antibodies (listed below). After 3×10min washes in TBST, the membrane was incubated at room temperature for additional two hours with secondary antibodies. The membrane was washed 3×10min in TBST and then visualized. Antibody concentrations for western blots were as follows: GFP (abcam ab290) 1:5000; β-actin (servicebio GB12001) 1:2000; His (proteintech 66006-1-Ig) 1:5000. anti-HA Ab and Anti-mouse IgG (H+L) were purchased from Proteintech.

### Statistics

Bar graphs with error bars were presented with mean and s.d. All of the experiments were conducted with independent *C. elegans* animals for an indicated N times. Statistical analysis was performed with the two-tailed Student’s t-test.

## Data availability

All related data and materials are available upon request.

## Acknowledgments

We are grateful to Dr. Guangshuo Ou’s lab for their technical supports and suggestions, and the members of the Guang lab for their comments. We are grateful to the International *C. elegans* Gene Knockout Consortium, and the National Bioresource Project for providing the strains. Some strains were provided by the CGC, which is funded by NIH Office of Research Infrastructure Programs (P40 OD010440). This work was supported by grants from the Chinese Ministry of Science and Technology (2017YFA0102903), the National Natural Science Foundation of China (Nos. 81501329, 31671346, 91640110, 31870812 and 31871300), the Major/Innovative Program of Development Foundation of Hefei Center for Physical Science and Technology (2017FXZY005), and CAS Interdisciplinary Innovation Team.

## Author contributions

C.Z. and S.G. designed the project; C.Z., C.W., X.W., Y.Y., W.L. D.X, M.H. S.L. and X.Z., performed research; C.Z., C.W. X.W. and Y.Y. contributed new reagents/analytic tools; C.Z. and C.W. analyzed data; C.Z. and S.G wrote the paper.

## Additional Information

The authors declare no competing financial interests.

## Supplementary figure legends

**Fig. S1. Construction of TOFU transgenes.** (A) Images of TOFU-1::GFP and TOFU-2::GFP in cut gonad. (B) Images of TOFU-6::GFP, GFP::PRG-1 and PGL-1::GFP in cut gonad. (C) Images of GFP::PRG-1 and the P-granule marker mRuby::PGL-1 in germline cells. (D) Bar diagram displaying brood size of indicated animals. Worms were grown at 20.

**Fig. S2. TOFU-6 is required for 21U-RNA biogenesis.** (A) Scatter plot comparing 21U-RNA cloning frequencies between wild type and *tofu-6* mutant. (B) Images of GFP::PRG-1 in indicated animals (top) and the relative GFP intensities were measured by ImageJ (bottom), ^***^ p<0.01.

**Fig. S3. PID-1 and PICS-1 are required for the granule formation of TOFU-6.** (A) Schematic of *pid-1* and *pics-1* alleles. (B) Images of TOFU-6::GFP in mitosis and meiosis regions in indicated animals.

**Fig. S4. PID-1 and PICS-1 are required for 21U-RNA biogenesis.** (A, B) Scatter plot comparing 21U-RNA cloning frequencies between wild type and *pid-1(ust64)* (A) and *pics-1(tm2417)* (B) mutant worms. (C) Images of PID-1::GFP in cut gonad. (D) Images of PICS-1::GFP in cut gonad. (E) Bar diagram displaying brood size of indicated animals. Worms were grown at 20.

**Fig. S5. PID-1 is required for the granule formation of PICS-1.** (A) Images of PICS-1::GFP after RNAi targeting indicated genes. (B) Western blotting of PICS-1::GFP after RNAi targeting indicated genes. (C) Images of PICS-1::GFP in mitosis and meiosis regions in indicated animals. (D) Images of PICS-1::GFP;*tofu-6(ust94)* after RNAi targeting *pid-1.*

**Fig. S6. Genetic requirements of the subcellular localization of TOST-1 and ERH-2.** (A, B) Images of TOST-1::GFP (A) and ERH-2::GFP (B) in cut gonad. (C) Images of ERH-2::GFP after RNAi targeting *tost-1*. (D) Western blotting of ERH-2::GFP after RNAi targeting *tost-1*. (E) Images of TOST-1::GFP in cut gonad t after RNAi targeting indicated genes. (F) Images of ERH-2::GFP after RNAi targeting indicated genes.

**Fig. S7. Summary of IP-MS results.** Top ten hits of TOFU-6, PICS-1, TOST-1 and ERH-2 are listed, respectively.

**Fig. S8. TOFU-6, PICS-1, ERH-2 and TOST-1 can accumulate in the nucleus.** (A, B) Images of TOFU-6::GFP, PICS-1::GFP, ERH-2::GFP and TOST-1::GFP in embryos (A) and in oocyte (B) after RNAi targeting indicated genes.

**Table S1:** IP-MS results of TOFU-6, TOST-1, PICS-1 and ERH-2.

**Table S2:** Strains used in the work.

**Table S3.** Primers used for transgene construction and sgRNA-mediated gene editing.

